# Circadian-Modulated Thresholds for Sleep Patterns in Aging and Narcolepsy

**DOI:** 10.64898/2026.08.02.742270

**Authors:** Chenggui Yao, Xiaoping Wu, Zijun Ning, Dongping Yang

## Abstract

In the two-process model of sleep-wake regulation, circadian-modulated thresholds time every sleep onset and awakening, yet they remain free parameters rather than quantities derived from neuronal dynamics. This limitation leaves the framework unable to predict how sleep patterns change when neuromodulatory drive is altered, as in aging and narcolepsy. Here we close that gap by deriving closed-form, circadian-modulated threshold expressions from the Phillips-Robinson model with explicit orexinergic excitation of the wake-promoting population. Within this single threshold geometry, aging and narcolepsy appear as opposite deformations along the orexinergic axis: age-related hyperexcitability of orexin neurons elevates the sleep-onset boundary and creates a fragility regime in which minor nocturnal disturbances trigger premature awakenings, whereas orexin loss depresses the same boundary toward the wake-onset threshold and produces the rapid state fragmentation of narcolepsy. Concurrently, reduced circadian amplitude compresses the inter-threshold corridor, advancing sleep onset and shortening sleep duration. These results convert the classical two-process thresholds from descriptive conveniences into mechanistic organizers of sleep–wake dynamics across healthy aging and orexin deficiency.

**Author summary:** The classical two-process model of sleep relies on a pair of switching thresholds that have been imposed by hand rather than derived from the neurons that actually stabilize sleep and wakefulness. Here we remove this limitation: starting from a biophysical mean-field model of the sleep- and wake-promoting populations, and adding the orexin system that stabilizes arousal, we derive the sleep-onset and awakening thresholds analytically from the bifurcation geometry of the underlying dynamical system. These closed-form thresholds depend explicitly on circadian phase, homeostatic state, and orexinergic tone, revealing that aging and narcolepsy are opposite deformations of a single threshold corridor along one orexinergic axis. In aging, orexin hyperexcitability raises the sleep-onset barrier past a sharp “arousal fragility boundary,” beyond which a minor disturbance triggers irreversible awakening; in narcolepsy, orexin loss collapses the same barrier and fragments sleep while paradoxically preserving total sleep time. We further find, contrary to common assumption, that orexin sustains wakefulness chiefly by raising the barrier to falling asleep rather than by resisting awakening. This work turns phenomenological sleep thresholds into physics-derived organizers of behavior, providing a mechanistic bridge from neuronal circuitry to whole-organism sleep dynamics.

## 1 Introduction

Sleep–wake timing is classically described by the two-process model, in which a homeostatic sleep pressure interacts with a circadian pacemaker to schedule sleep onset and offset [1, 2]. The model’s predictive power rests on a pair of thresholds that bound the homeostatic drive: sleep begins when pressure rises to the upper boundary, and wake resumes when it falls to the lower one. Yet these thresholds are imposed phenomenologically. They are not derived from the neuronal circuitry that actually stabilizes behavioral state, and therefore cannot explain how sleep consolidation changes when that circuitry is altered by age or disease [3, 4].

The Phillips-Robinson (PR) model advanced this picture by embedding two-process logic in a mutual-inhibition dynamical system between wake- and sleep-promoting nuclei [5– 8]. State transitions then emerge from physiologically interpretable, time-varying thresholds rather than from an externally prescribed schedule. Even so, the PR thresholds remain fixed by model parameters and do not incorporate the orexin (hypocretin) system—the neuromodulatory drive now recognized as the principal stabilizer of wakefulness [9, 10]. Without an explicit orexinergic term, the framework cannot represent how firmly the system is held in wake or sleep, nor how that holding force varies across individuals, ages, and pathologies.

Orexin warrants this treatment because it occupies a uniquely decisive position in the sleep-wake control. Orexinergic neurons of the lateral hypothalamus excite monoaminergic and cholinergic arousal nuclei, consolidating wakefulness and suppressing premature transitions into sleep. The clinical significance of this role is revealed at two extremes. Selective loss of orexin-producing neurons abolishes the drive and yields narcolepsy type 1, with fragmented vigilance, intrusive sleep episodes, and collapse of stable wake-sleep boundaries [11–14]. Aging, by contrast, is not a simple reduction of orexinergic signaling: surviving hypocretin/orexin neurons become hyperexcitable, and this hyperexcitability–rather than cell loss alone–contributes to the lighter, more fragmented, and phase-advanced sleep of older adults [15, 16]. Orexin is therefore the physiological quantity that the two-process model leaves implicit and that standard PR thresholds fix by hand: its deficiency and its hyperexcitable excess should perturb the same switching boundaries in opposite directions.

Here we reduce an extended PR framework with explicit orexinergic excitation of the wake-promoting population [5, 6] and obtain the sleep and wake thresholds analytically by linear stability analysis. The resulting closed-form expressions are circadian-modulated and orexin-dependent, so that changes in circadian amplitude, homeostatic capacity, and orexinergic tone map directly onto deformations of threshold geometry. We use this geometry to address two problems usually treated in isolation. First, we analyze aging by varying circadian drive, homeostatic drive, and orexinergic tone independently, reproducing advanced timing, reduced consolidation, and increased nocturnal fragility [16, 17]; in particular, orexinergic hyperexcitability elevates the sleep-onset boundary into a regime where small disturbances can trigger irreversible awakenings. Second, we account for narcolepsy as orexin deficiency that lowers the sleep-onset boundary toward the wake-onset threshold, thereby destabilizing wakefulness and facilitating sleep intrusion [11, 12]. In both cases, orexin-regulated and circadian-modulated thresholds supply a common quantitative language for sleep patterns in healthy aging and orexin deficiency.

## 2 Results

### 2.1 An extended sleep–wake model with explicit orexinergic drive

Our analysis is grounded in a mean-field population model that extends the Phillips–Robinson (PR) framework by incorporating orexin (Orx) as an explicit wake-promoting population rather than subsuming it into a lumped monoaminergic system [5, 6, 18]. Drawing on the sleep model of Fulcher *et al*. [14] and informed by the well-characterized neuroanatomical connectivity [9], our formulation retains the essential flip-flop architecture while adopting a more complete set of inter-population pathways. The model comprises three interacting neural populations: the sleep-promoting ventrolateral preoptic area (VLPO), an aggregate non-Orx monoaminergic wake-promoting population (MA), and Orx, together with circadian and homeostatic drives. The network architecture is defined by three classes of interaction. (i) VLPO and MA mutually inhibit each other, in accord with the canonical flip-flop organization of sleep–wake control [19–22]. (ii) Orx and MA form a sign-asymmetric reciprocal pair: Orx provides dense excitatory drive that tonically supports wake-active monoaminergic neurons [23–25], whereas serotonin and noradrenaline released by MA inhibit Orx through *α*_2_-adrenergic and 5-HT_1A_ receptors coupled to GIRK channels [26–28]. (iii) VLPO additionally provides a direct inhibitory projection to Orx [29, 30]; this pathway silences Orx during sleep and complements the VLPO–MA flip-flop. A comprehensive review of these interactions is given in Ref. [9].

Sleep-wake transitions are further regulated by circadian and homeostatic processes. The circadian drive *C* represents SCN-derived rhythmic input entrained by the light-dark cycle [31, 32]. In the model, this drive suppresses VLPO activity and promotes LC-mediated arousal, consistent with hypothalamic circadian output pathways involving the dorsomedial hypothalamus [20, 33]. The homeostatic drive *H* represents sleep pressure that accumulates during wakefulness and dissipates during sleep, in accordance with the two-process framework [1–4]. Physiologically, this process reflects the buildup of sleep-promoting substances such as adenosine, which facilitate VLPO recruitment and bias the system toward sleep [34–37].

A schematic of the model architecture is shown in Fig. A1 (Model and Methods); the full dynamical equations and all parameter values are provided in Appendix A and Table 1. Parameters are chosen to produce a stable baseline state characterized by regular sleep-wake cycling with sharp state transitions. Below we derive the corresponding orexin-regulated and circadian-modulated thresholds and show how their geometry accounts for sleep patterns in aging and narcolepsy.

**Table 1.** Nominal parameter values. Here, *ν*_*ab*_ denotes the effective coupling from population *b* to population *a*.

| Parameter | Value | Unit | Description |
| --- | --- | --- | --- |
| $\nu_{mv}$ | -1.8 | mV s | Inhibitory coupling from VLPO to MA |
| $\nu_{mx}$ | 0.1–0.7 | mV s | Effective excitatory coupling from Orx to MA |
| $\nu_{vm}$ | -1.8 | mV s | Inhibitory coupling from MA to VLPO |
| $\nu_{xm}$ | -0.3 | mV s | Effective excitatory coupling from MA to Orx |
| $\nu_{xv}$ | -1.8 | mV s | Inhibitory coupling from VLPO to Orx |
| $\nu_{vc}$ | -0.6 | mV | Circadian input to VLPO |
| $\nu_{xc}$ | 2.0 | mV | Circadian input to Orx |
| $\nu_{vh}$ | 0.7 | mV nM<sup>-1</sup> | Homeostatic input to VLPO |
| $\tau_m$ | 10 | s | Time constant of MA population |
| $\tau_v$ | 10 | s | Time constant of VLPO population |
| $\tau_x$ | 120 | s | Time constant of Orx population |
| $A_m$ | 1.0 | mV | Input from groups including acetylcholine and orexin |
| $A_v$ | -1.0 | mV | The time-averaged inputs to each population |
| $A_x$ | 1.0 | mV | Constant background input to Orx |
| $\tau_{hm}$ | 25 | h | Time constant for homeostatic buildup during wake |
| $\tau_{hv}$ | 7 | h | Time constant for homeostatic decay during sleep |
| $\mu$ | 13 | nM | Maximum production level of homeostatic drive |
| $\eta$ | 2.3 | s<sup>-2</sup> | Half-saturation parameter for homeostatic production |
| $Q_{\max}$ | 100 | s<sup>-1</sup> | Maximum firing rate |
| $\theta$ | 10 | mV | Mean firing threshold relative to rest |
| $\sigma'$ | 3 | mV | Standard deviation of firing threshold |
| $V_{th}$ | -2.0 | mV | Wake threshold for homeostatic switching |

### 2.2 Aging sleep-wake dynamics as a probe of circadian-modulated thresholds

Advancing into the fifth and sixth decades of life, human sleep undergoes a collection of well-characterized architectural changes: (1) sleep timing advances, with both bedtimes and rise times shifting earlier relative to younger adults; (2) total sleep duration shortens; (3) sleep becomes more fragmented, exhibiting more frequent awakening, arousal, and transitions to lighter stages [16]. These features are typically listed as separate empirical observations, yet in the threshold framework they share a common mechanistic reading. Sleep timing is set by the crossings of homeostatic pressure with the switching thresholds; sleep duration is the interval between those crossings; and fragmentation reflects how easily a perturbation can push the system across a nearby boundary. All three hallmarks of aging sleep can therefore be read as signatures of alter threshold geometry.

We first establish a young-adult baseline. Using the parameter-selection framework described in Ref. [38], we obtain physiologically realistic sleep-wake transitions (Fig. 1a). These transitions are understood through fast-slow reductions, in which bifurcations of the fast subsystem are driven by two aggregated slow variables, *D*_*x*_ and *D*_*v*_ [7, 14, 39]. The resulting phase diagram in the (*D*_*x*_, *D*_*v*_) plane (Fig. 1b) is partitioned into wake, sleep, and bistable regions by two critical boundaries (*f*_1_ and *f*_2_) whose positions depend on orexinergic drive. Under circadian and homeostatic forcing, the trajectory evolves slowly through this plane and switches whenever it crosses a boundary, thereby generating a complete sleep-wake cycle.

**Fig. 1.**
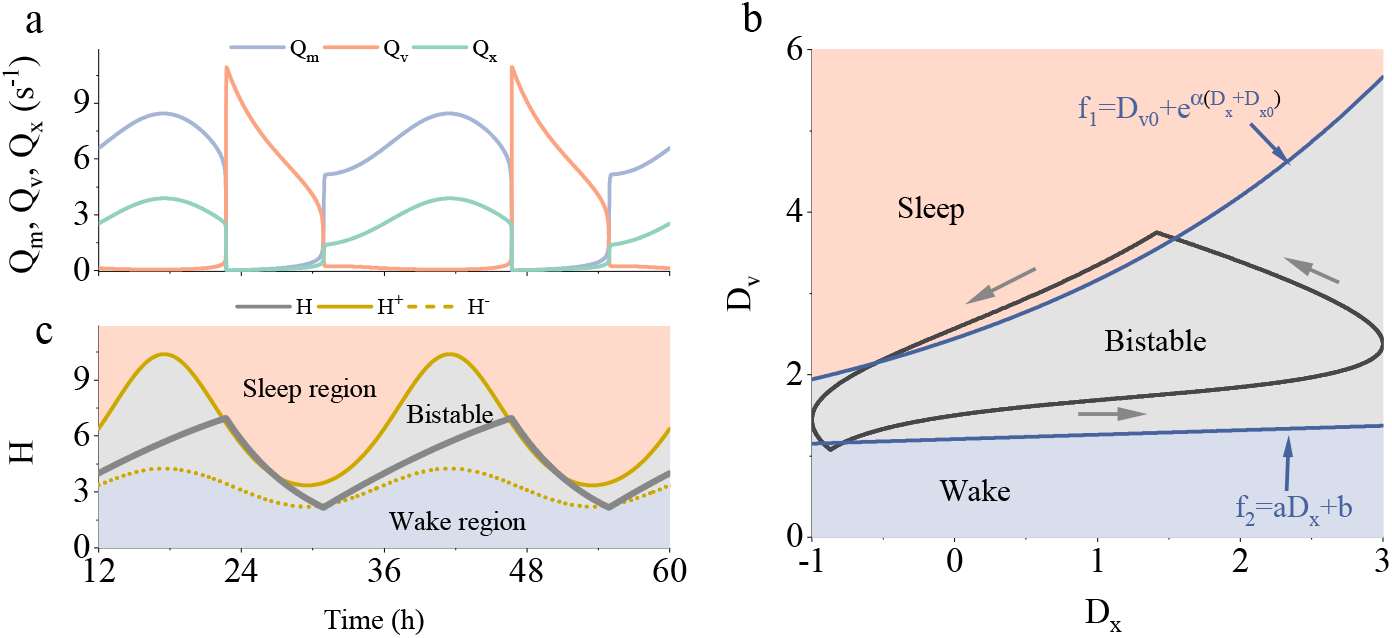
Baseline sleep-wake cycle and circadian-modulated thresholds. **(a)** Representative baseline sleep-wake cycle of a young adult. Time series of the wake-active population firing rate *Q*_*m*_, sleep-active population firing rate *Q*_*v*_ , and orexinergic population firing rate *Q*_*x*_. **(b)** Dynamical phase diagram in the (*D*_*x*_, *D*_*v*_ ) plane, partitioned into wake, sleep, and bistable regions. The nominal trajectory evolves slowly under circadian and homeostatic forcing and generates a full sleep-wake cycle by crossing the two critical boundaries. The sleep-onset boundary can fitted by 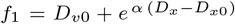, and the awakening boundary by *f*_2_ = *a D*_*x*_ + *b*. **(c)** Threshold-based switching mechanism. Homeostatic sleep pressure *H* (gray line) is plotted together with the analytically derived sleep-onset threshold *H*^+^ (solid orange line) and awakening threshold *H*^−^ (dashed orange line). Sleep is initiated when *H* crosses above *H*^+^ and terminated when *H* falls below *H*^−^.

To connect this geometry to observable physiology, we separate the homeostatic variable *H* from the circadian signal *C*, and analytically transform *f*_1_ and *f*_2_ into time-dependent thresholds for homeostatic sleep pressure: a sleep-onset threshold *H*^+^(*t*) and an awakening threshold *H*^−^(*t*) (Eq. 16, see Appendix B for the full derivation). Fig. 1c illustrates the resulting switching rule. Sleep begins when *H* (gray line) rises above *H*^+^ (solid orange line), and awakening occurs when *H* falls below *H*^−^ (dash-dotted orange line). Circadian modulation therefore continuously reshapes the homeostatic conditions required for each transition, linking the underlying drives to the timing of sleep onset and offset.

Having established the threshold framework, we next ask how age-related loss of circadian amplitude reshapes sleep timing and duration. Figure 2a shows *H*^+^(*t*) and *H*^−^(*t*) for circadian amplitude ratios *r* = 1.0, 0.70, and 0.42. Reducing *r* affects the two thresholds asymmetrically: *H*^+^ drops substantially, while *H*^−^ decreases only modestly. The resulting compression of the bistable corridor advances both sleep onset and wake onset, but the wake advance is larger, so total sleep duration shortens slightly (Fig. 2c, upper panel). These predictions match the earlier timing and modest shortening of sleep in older adults [16, 17].

**Fig. 2.**
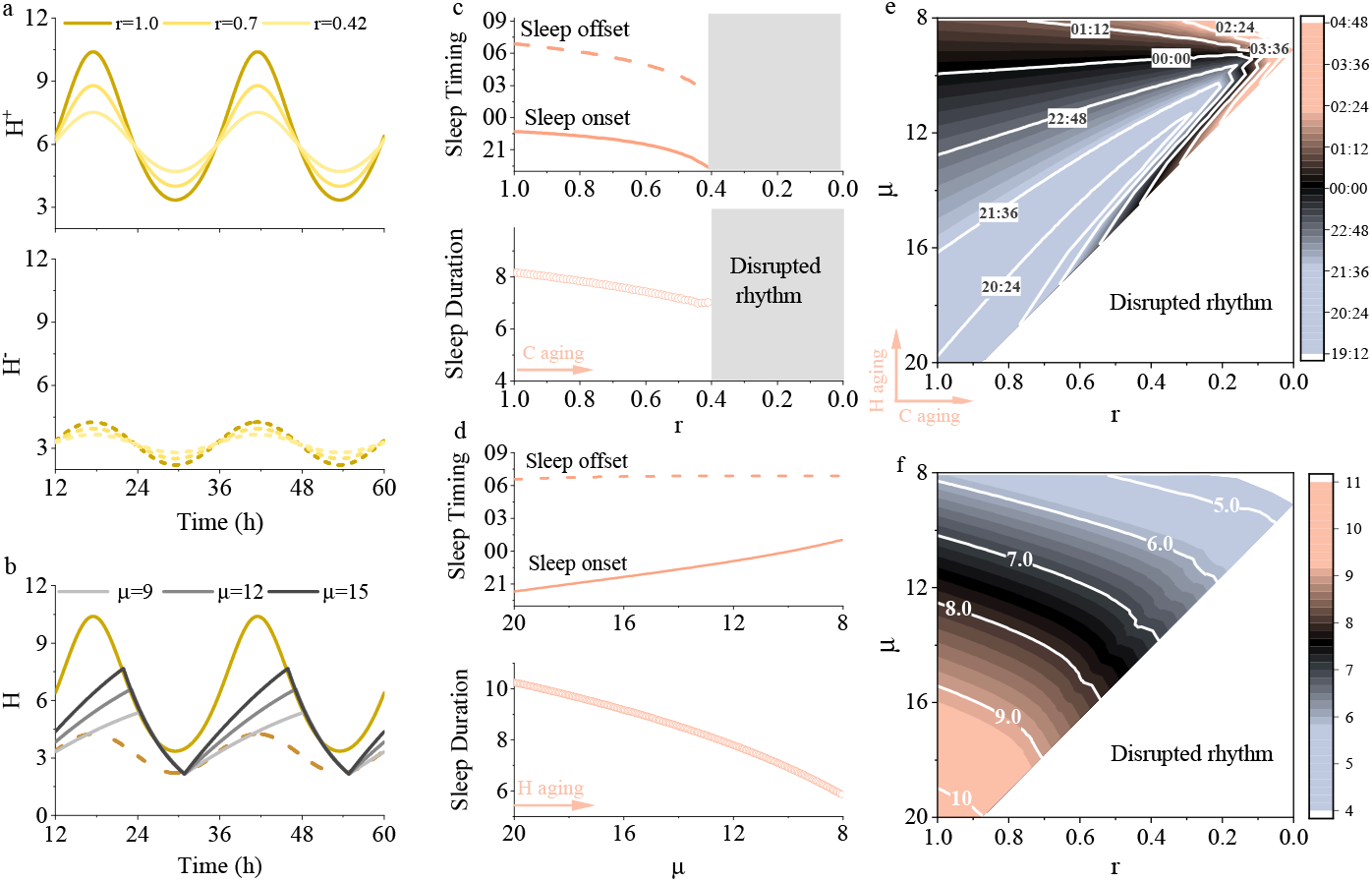
Aging sleep-wake dynamics as a probe of circadian-modulated thresholds. **(a)** Effect of reduced circadian amplitude on the thresholds. Time series of *H*^+^(*t*) and *H*^−^(*t*) for *r* = 1.0, 0.70, and 0.42, where *r* represents the ratio of aged to baseline circadian amplitude. Decreasing circadian amplitude compresses the vertical separation between the two thresholds. **(b)** Effect of reduced homeostatic maximum on sleep pressure dynamics. Time series of *H*(*t*) for *µ* = 9, 12, and 15, where *µ* parameterizes the maximum attainable homeostatic pressure. Lower values of *µ* reduce the peak level of *H* accumulated during wakefulness. **(c)-(d)** Dependence of sleep characteristics on aging parameters *r* and *µ*, respectively. Top panels: sleep timing (onset and offset); bottom panels: total sleep duration per cycle. **(e)-(f)** Contour plots for sleep onset and sleep duration as functions of *µ* and *r*.

Figure 2b examines a complementary pathway by lowering the homeostatic maximum *µ*. As *µ* decreases from 15 to 9, peak *H* falls so that *H* takes longer to reach *H*^+^ and sleep onset is delayed. Wake-onset time remains nearly unchanged (Fig. 2d, upper panel), as the lower peak is partly offset by a faster approach to *H*^−^. The net result is a clearer reduction in total sleep duration than that produced by circadian attenuation alone (Fig. 2d, lower panel). The two pathways therefore erode sleep through distinct geometries: circadian amplitude reduction compresses the threshold gap and shifts the sleep window earlier, whereas homeostatic weakening truncates sleep mainly by delaying its onset.

Their joint action is quantified in Figs. 2e and 2f. Reducing *r* alone advances onset and mildly shortens sleep; reducing *µ* alone delays onset and more strongly curtails duration. Along the diagonal these effects partially cancel, so timing can remain nearly stationary despite substantial degradation of both subsystems. Blank regions–where stable cyclic behavior disappears–concentrate at low *r* with moderate-to-high *µ*, indicating that severe circadian amplitude loss can precipitate arrhythmia, while a concurrently weakened homeostatic drive postpones this destabilization by reducing the pressure excursion that must fit within the narrowed corridor. Homeostatic aging thus acts as a partial buffer against circadian-driven fragmentation, an interaction invisible when each pathway is varied alone.

### 2.3 Orexinergic hyperexcitability as a mechanism of age-related sleep fragility

Circadian attenuation and homeostatic weakening reshape the global architecture of sleep by compressing or shifting the threshold corridor. We now turn to a third aging axis: the orexinergic system. Aging-related neurodegeneration in the lateral hypothalamus does not simply silence orexin neurons; the surviving population becomes hyperexcitable [15]. We capture this pathophysiology by taking the maximum orexin firing rate evoked by excitatory drive *A*_*x*_, denoted as 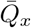, as a single varying parameter, and investigate how its elevation reshapes both the baseline sleep-wake cycle and the vulnerability of sleep to external perturbation.

The most consequential effect of orexinergic hyperexcitability is microscopic: it erodes the resilience of sleep against transient disturbances. We applied a five-minute impulsive stimulus (transient increase Δ*A*_*m*_) to the MA population during consolidated sleep and tracked the ensuing dynamics at 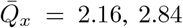, and 3.86 with sleep drive fixed at *D*_*v*_ = 2.0 (Fig. 3a). At the low excitability 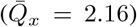, the stimulus elicits only a fleeting arousal and sleep resumes within minutes (Fig. 3a, top). At the intermediate one 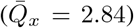, the same stimulus triggers a prolonged waking episode exceeding one hour before sleep returns (Fig. 3a, middle). At the high excitability 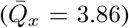, a single transient stimulus ejects the system from sleep permanently for the remainder of the circadian night (Fig. 3a, bottom).

**Fig. 3.**
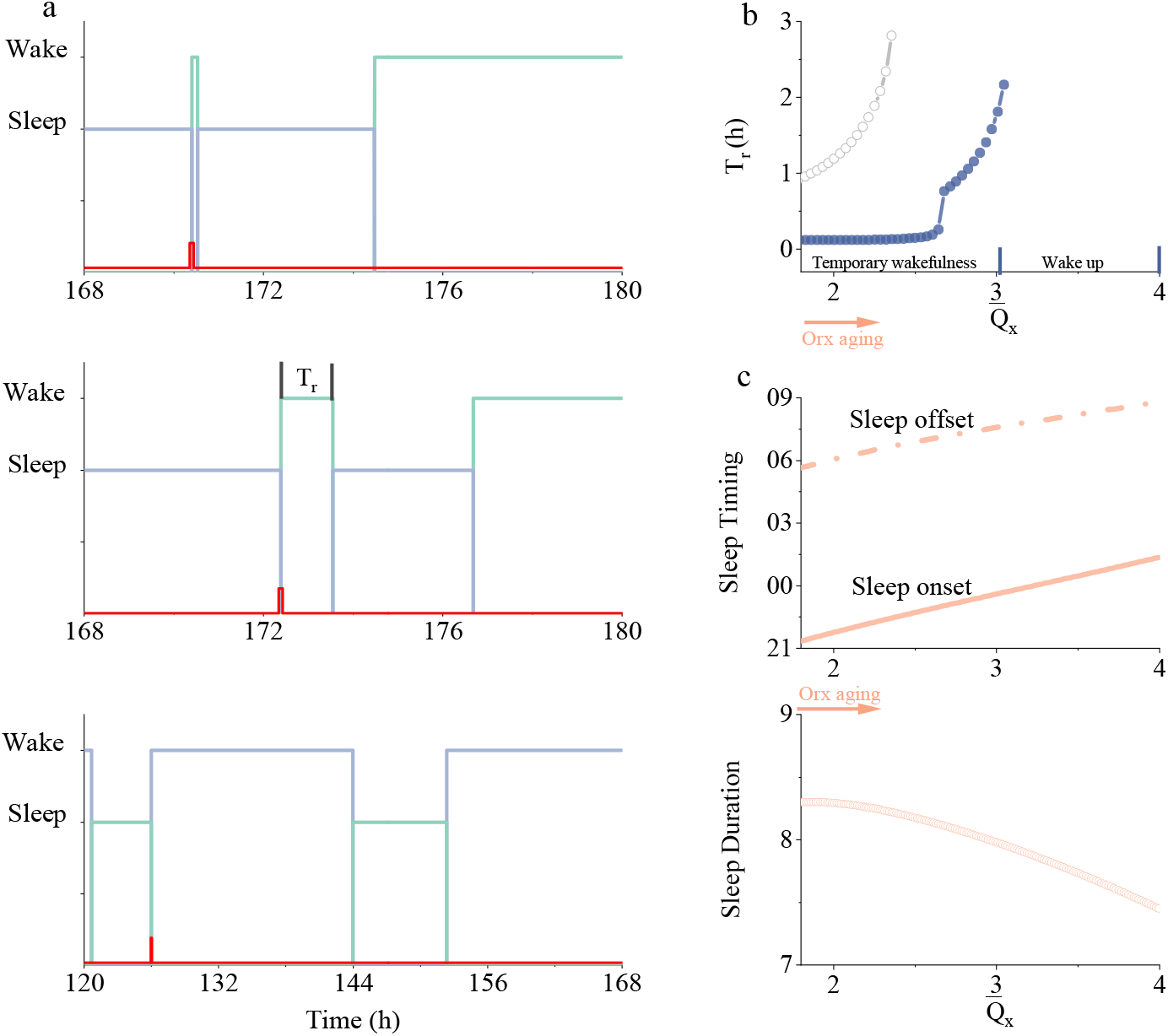
Orexinergic hyperexcitability erodes sleep resilience and defines an arousal fragility boundary. **(a)** Responses to a brief external stimulus (a transient pulse Δ*A*_*m*_, the red line, applied during sleep) for three levels of orexinergic excitability (from top to bottom: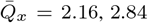, and 3.86). **(b)** Duration of stimulus-evoked wakefulness as a function of 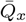, demonstrating a divergence at a critical excitability beyond which small transient perturbation can lead to irreversible awakening. **(c)** Sleep timing (onset and offset, upper) and the total sleep duration (bottom) as a function of 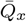.

Figure 3b distils this progression into a single curve by plotting stimulus-evoked wake duration (*T*_*r*_, as shown in Fig. 3a) as a function of 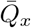. The duration rises gradually at first, then steepens sharply and diverges at a critical excitability, beyond which any transient perturbation produces irreversible awakening. This critical point defines an *arousal fragility boundary*: below it, sleep can absorb disturbances and self-repair; above it, even mild perturbations shatter sleep continuity irrecoverably. Such a sharp boundary offers a compact mechanistic account of why nocturnal awakenings in aging populations not only become more frequent but increasingly fail to resolve back into consolidated sleep [16].

The same orexinergic tone also reshapes the baseline cycle (Fig. 3c). As 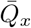 increases, both sleep onset and offset are delayed, but onset shifts more than offset, so total sleep duration contracts monotonically. As shown next, this asymmetry reflects the differential sensitivity of the two thresholds to orexinergic tone–the same mechanism that underlies the graded fragility above.

The dynamical origin of these brief awakenings is the saddle-node structure of the fast subsystem (Eq. 7, Fig. 4a): increasing Δ*A*_*m*_ annihilates the sleep fixed point. The corresponding hysteresis loops in the (Δ*A*_*m*_, *D*_*v*_) plane (Fig. 4b) show that higher 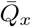 progressively inflates the *D*_*v*_–equivalently, the homeostatic pressure *H*–required to return to sleep, geometrically widening the wake-trapped corridor. Orexinergic aging therefore does not merely make awakening easier; it makes falling back asleep disproportionately harder.

**Fig. 4.**
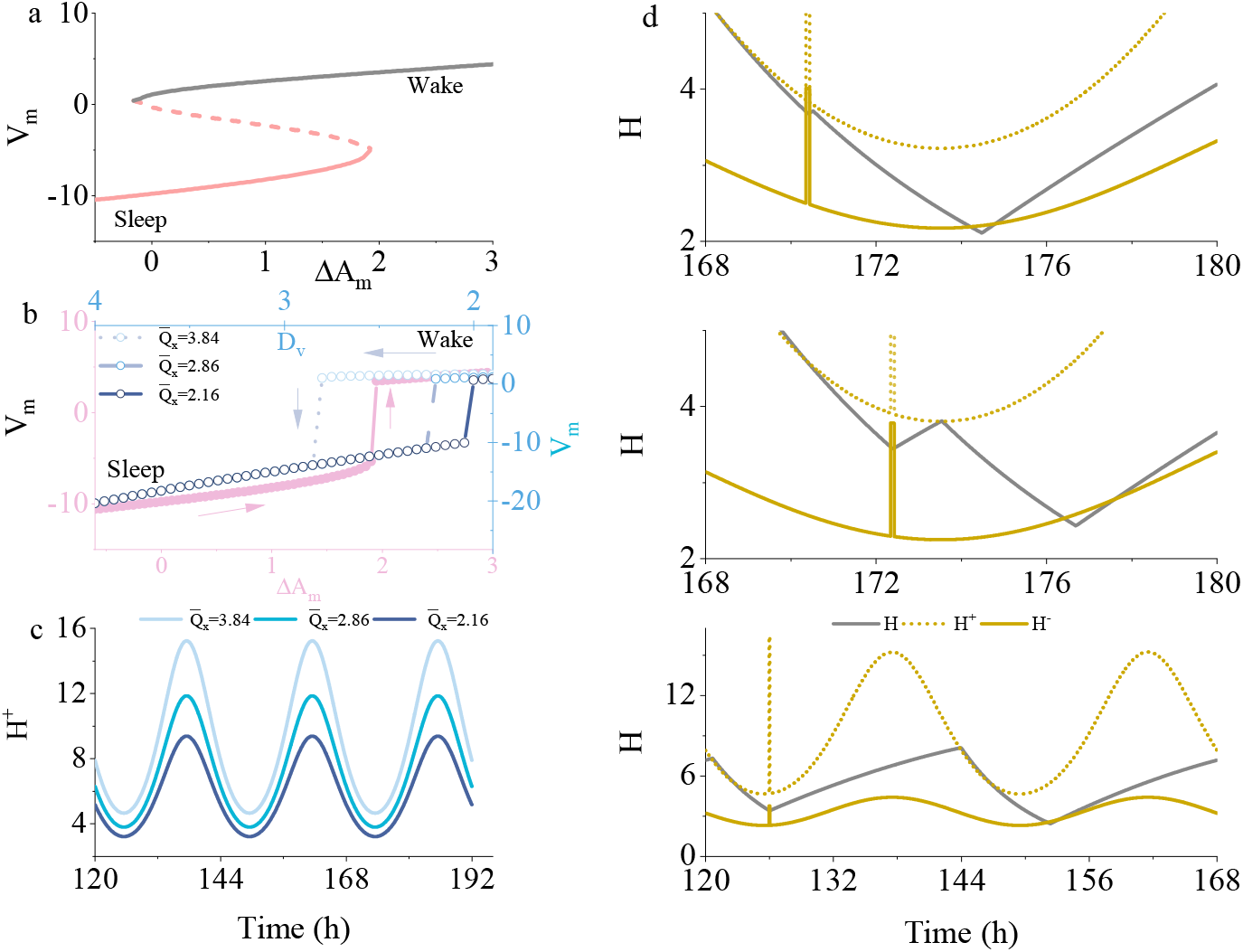
Threshold mechanism of aging-related orexinergic hyperexcitability. **(a)** Saddle-node bifurcation diagram as a function of stimulus amplitude Δ*A*_*m*_, illustrating the critical stimulus strength required to trigger a sleep-to-wake transition. **(b)** Hysteresis loops in the (Δ*A*_*m*_, *D*_*v*_ ) plane. Increasing Δ*A*_*m*_ drives the system from sleep to wake, whereas increasing *D*_*v*_ drives the reverse transition. The three curves correspond to 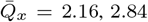, and 3.86, showing that higher orexinergic excitability widens the wake-trapped corridor of the hysteresis region. **(c)** Effect of orexinergic excitability (aging) on the circadian sleep thresholds. Time series of *H*^+^ for 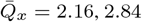, and 3.86. Increasing orexinergic excitability elevates the sleep-onset threshold. **(d)** Homeostatic sleep pressure *H* together with *H*^+^ and *H*^−^ after a brief external stimulus for three levels of orexinergic excitability (from top to bottom: 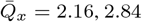, and 3.86).

Recast in threshold language, increasing 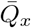 elevates *H*^+^ substantially while leaving *H*^−^ largely unchanged (Fig. 4c; Appendix B). This asymmetric shift narrows the corridor traversed by *H* during sleep and governs both baseline timing and resilience to perturbation. Figure 4d makes the graded fragility explicit. The brief pulse transiently elevates *H*^−^, opending a momentary gateway through which declining *H* escapes into the wake-permissive domain. Return to sleep, however, is controlled by *H*^+^, which rises with 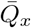. At low 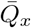 , accumulating *H* reaches *H*^+^ quickly and sleep resumes. At intermediate 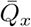, the elevated *H*^+^ demands a longer wake sojourn before sufficient pressure accumulates. At high 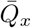 , *H* cannot reach *H*^+^ within any physiologically plausible interval, and the system remains irreversibly trapped in wake.

### 2.4 Narcolepsy as a probe of orexin-modulated thresholds

Whereas the preceding analysis examined orexinergic hyperexcitability–a hallmark of aging– we now consider the opposite pathological extreme: the progressive loss of orexinergic tone, the defining lesion of narcolepsy type 1 [11–13]. In the model, orexin’s influence on the wake-promoting MA population is set by the coupling strength *ν*_*mx*_; reducing this parameter mimics the graded degeneration of orexin-producing neurons observed clinically [13].

Figure 5a illustrates the essential dynamical consequence of orexin deficiency. Rather than a single consolidated sleep bout and a single consolidated wake bout per cycle, the system fragments into rapid, involuntary oscillations between the two states. In a healthy regime, *H*^+^ and *H*^−^ are well separated, creating a wide bistable corridor that prevents spontaneous sleep intrusions and unwanted awakenings. When *ν*_*mx*_ is reduced, that gap collapses. During wakefulness, the lowered *H*^+^ is reached after only a modest accumulation of homeostatic pressure, tipping the system into sleep prematurely. Once asleep, *H* need fall only slightly before encountering *H*^−^, triggering re-awakening almost immediately. Each state is therefore abandoned nearly as soon as it is entered.

**Fig. 5.**
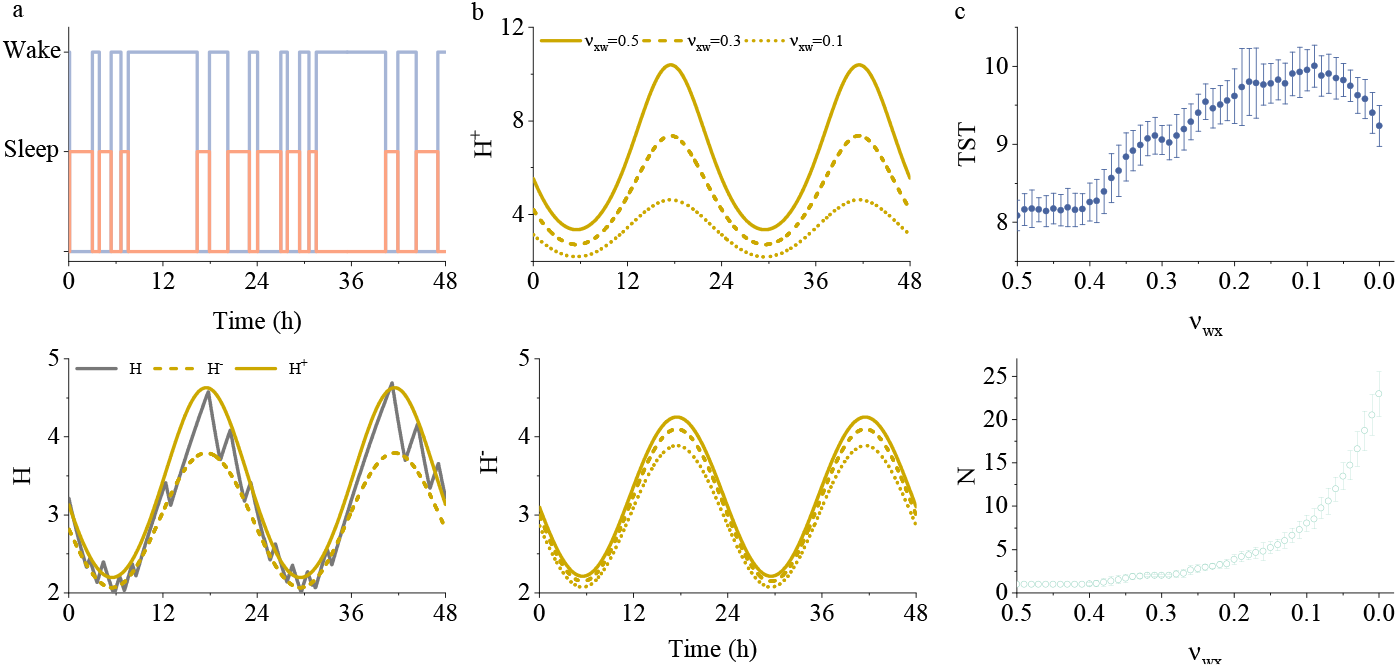
Orexin deficiency destabilizes the sleep-wake flip-flop and produces narcoleptic dynamics. **(a)** Schematic of narcolepsy induced by reduced orexinergic tone (*ν*_*mx*_ = 0.1, upper panel). The corresponding time series of homeostatic sleep pressure *H*, together with *H*^+^ and *H*^−^, are shown in the lower panel; the diminished separation between *H*^+^ and *H*^−^ permits rapid, involuntary oscillations between sleep and wake. **(b)** *H*^+^ (upper) and *H*^−^ (lower) over the circadian cycle for *ν*_*mx*_ = 0.5 (intact), 0.3 (moderate loss), and 0.1 (severe loss). Progressive reduction of *ν*_*mx*_ compresses the hysteresis gap between the two thresholds. **(c)** Total sleep time per 24-hour period (TST; upper) and mean number of sleep-wake transitions per day (lower) as functions of *ν*_*mx*_. As *ν*_*mx*_ decreases, TST increases modestly while the transition count rises sharply.

This compression is quantified in Fig. 5b, for *ν*_*mx*_ = 0.5, 0.3, and 0.1. Both thresholds decrease as *ν*_*mx*_ falls, but at different rates: *H*^+^ drops precipitously, whereas *H*^−^ decreases only modestly. It is this asymmetry–the disproportionately steep fall of *H*^+^–that compresses the bistable corridor from above and produces state fragmentation.

This result challenges the intuition that orexin stabilizes wakefulness primarily by raising the awakening threshold *H*^−^. In this model, the dominant effect of orexinergic tone is instead to elevate the *sleep-onset* threshold *H*^+^: orexin keeps the brain awake mainly by raising the barrier that homeostatic pressure must surmount before sleep can begin. When that barrier is removed by orexin loss, *H*^+^ collapses toward *H*^−^, and the wide hysteresis gap that normally prevents involuntary sleep intrusions during the biological day disappears.

Fig. 5c summarizes the population-level consequences. As *ν*_*mx*_ decreases from its healthy value, total sleep time per 24-hour period (TST) increases modestly–consistent with clinical observations that narcoleptic patients spend slightly more, not less, time asleep that controls [40]–while the daily transition count rises steeply below a critical range. The dissociation between preserved sleep quantity and shattered architecture is thus a natural consequence of threshold compression within the flip-flop framework.

Taken together with the aging results of § 2.2–2.3, these findings reveal a duality alonge the orexinergic axis. Orexinergic hyperexcitability (elevated 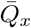) inflates the hysteresis gap, stabilizing wakefulness at the expense of sleep maintenance and rendering nocturnal sleep vulnerable to irreversible disruption by minor perturbations (Fig. 3 and 4). Orexin deficiency (reduced *ν*_*mx*_), by contrast, compresses the same gap, destabilizing both states and producing the rapid-cycling fragmentation of narcolepsy (Fig. 5). The healthy brain thus occupies a narrow operating regime in which orexinergic tone is calibrated to keep the hysteresis corridor wide enough for state consolidation yet flexible enough for timely transitions–a balance disrupted, in opposite directions, by aging and by narcoleptic neurodegeneration.

## 3 Discussion

The two-process model has long organized sleep research, yet its switching thresholds have remained phenomenological parameters rather than derived quantities. Starting from a neuronal population model with sleep-promoting, wake-promoting, and orexinergic populations, we obtained closed-form expressions for the sleep-onset threshold *H*^+^(*t*) and the awakening threshold *H*^−^(*t*) as explicit functions of circadian phase, homeostatic state, and orexinergic drive. Timing, duration, consolidation, and fragmentation then appear as geometric properties of a single circadian-modulated threshold corridor. Aging and narcolepsy deform that corridor along opposite directions of the orexinergic axis, while circadian and homeostatic aging provide complementary deformations. Attenuated circadian amplitude compresses the inter-threshold gap and advances the sleep window; reduced homeostatic capacity delays onset and further shortens sleep; orexinergic hyperexcitability elevates *H*^+^ into a fragility regime in which minor nocturnal disturbances can trigger irreversible awakening. Narcolepsy type 1 is the geometric dual: progressive orexin loss collapses the hysteresis gap mainly by dropping *H*^+^ toward *H*^−^, trapping the system in rapid-cycling fragmentation while preserving or slightly increasing total sleep time. In both cases, orexin acts principally by setting the height of the sleep-onset barrier rather than by merely deepening resistance to awakening.

These results matter for applications that already treat thresholds as the interface between physiology and behavior. Sleep-sufficiency metrics for shift workers [41], wearable model-guided interventions [42], threshold-referenced phenotyping of endogenous versus environmental control [43], and threshold-based readings of subjective sleepiness [44] all rely on the position of homeostatic drive relative to switching boundaries. As those boundaries have historically been phenomenological or fixed PR parameters, such models can adjust for circadian phase and homeostatic load but not for orexinergic variation in state-holding capacity. The present expressions supply that missing dimension: individual differences in circadian amplitude, homeostatic capacity, and orexinergic tone map onto identifiable deformations of *H*^+^ and *H*^−^, offering a mechanistic basis for personalized fatigue prediction and for evaluating whether a countermeasure acts via circadian, homeostatic, or orexinergic pathways.

Several simplifications constrain the present framework. The model treats each neural population as a mean-field ensemble and does not distinguish NREM from REM sleep; Incorporating REM-generating circuitry [45, 46] would be needed to capture stage architecture and, in particular, sleep-onset REM periods in narcolepsy. Orexinergic influence is parameterized by two scalars (*ν*_*mx*_ and 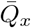), whereas the biological system comprises two peptides and two receptor subtypes with distinct targets [9]; a finer representation would enable subtype-selective pharmacology such as dual orexin-receptor antagonists [47]. Finally, the homeostatic process is modeled as a single exponential variable, whereas multi-timescale sleep debt my be required for chronic restriction. Despite these limits, the closed-form dependence of *H*^+^ and *H*^−^ on circadian amplitude, homeostatic capacity, and orexinergic tone provides a natural parameterization for wearable-derived individual profiles and for testing whether intermediate orexinergic phenotypes populate a continuous spectrum of state stability across aging cohorts.

**Fig. A1.**
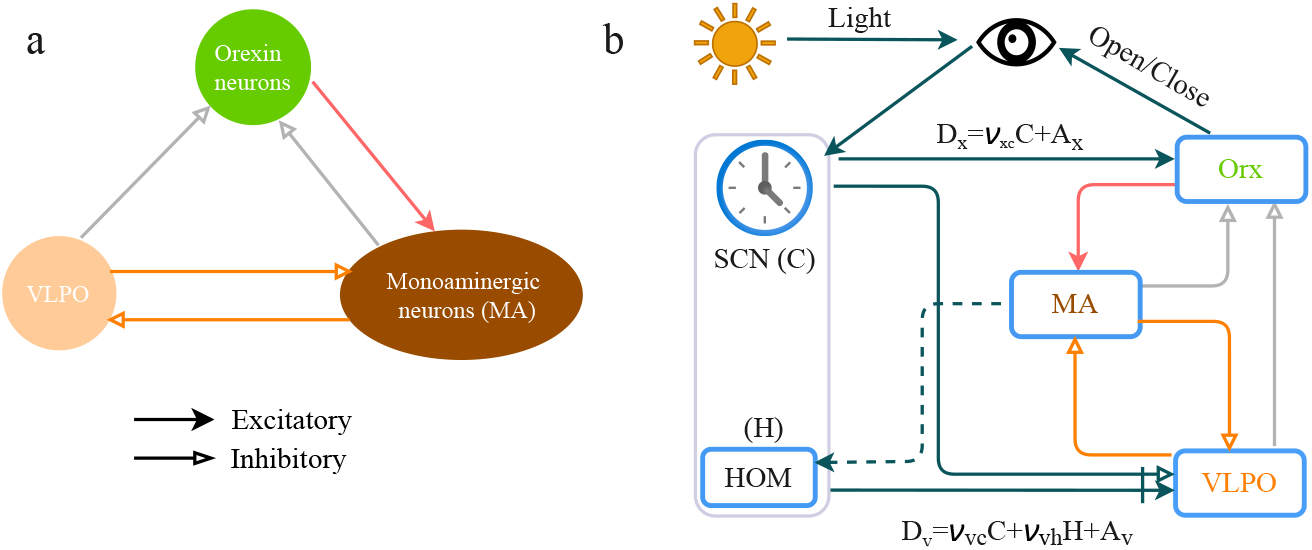
Schematic of the extended Phillips-Robinson sleep-wake model [6]. (a) Three mutually interacting neural populations: the sleep-active ventrolateral preoptic area (VLPO), an aggregate monoaminergic wake-promoting population (MA), and the orexinergic neurons (Orx) of the lateral hypothalamic area [9]. Reciprocal VLPO-MA inhibition implements the canonical sleep-wake flip-flop, while Orx provides a stabilizing excitatory drive to MA and receives inhibitory feedback from VLPO. (b) The circuit is coupled to the two-process model of sleep regulation: the circadian drive *C* from the suprachiasmatic nucleus (SCN), and the homeostatic drive *H*, which accumulates during wake and dissipates during sleep, jointly modulate VLPO activity. Arrows denote excitatory couplings; flat-ended bars denote inhibitory couplings

In summary, orexin-dependent circadian-modulated thresholds convert the classical two-process boundaries from descriptive conveniences into mechanistic organizers of sleep-wake dynamics. Aging and narcolepsy emerge as opposite deformations of one threshold geometry, and the healthy operating regime is the narrow corridor between them.

## 4 Model and Methods

### 4.1 Model

### 4.2 Mathematical formulation

The model architecture is summarized in Fig. A1. *V*_*a*_ (*a* = *m, v, x*) denotes the mean membrane potential of the MA, VLPO, and Orx populations, respectively, and *Q*_*a*_ denotes the corresponding firing rate. The model is presented as

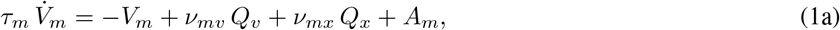

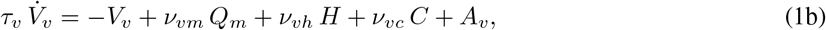

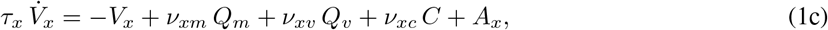

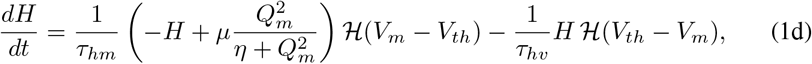

where *τ*_*a*_ are characteristic time constants, *A*_*a*_ are constant background inputs, and *ν*_*ab*_ is the effective coupling weight from population *b* to population *a*. The population firing rate obeys the sigmoidal transfer function

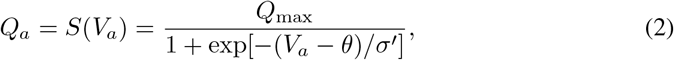

with *Q*_max_ the maximum firing rate, *θ* the mean firing threshold, and 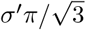 the standard deviation of firing thresholds across the population [6].

The signs of the coupling constants encode the architecture defined above: *ν*_*mv*_ *<* 0 and *ν*_*vm*_ *<* 0 enforce the MA-VLPO mutual inhibition; *ν*_*xv*_ *<* 0 and *ν*_*xm*_ *<* 0 implement the inhibition of Orx by VLPO and MA; and *ν*_*mx*_ *>* 0 captures the excitatory Orx → MA projection. The dimensionless circadian drive *C* originates from the SCN and is entrained to an effective period of ~ 24.15 h via retinal photic input [48]; it inhibits sleep-promoting neurons (*ν*_*vc*_ *<* 0) [20, 49] and excites orexinergic neurons (*ν*_*xc*_ *>* 0) [20, 50]. For analytical tractability, the circadian drive is modeled as a sinusoidal signal, *C*(*t*) = sin(*ω*_*c*_*t*) with 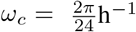, consistent with prior theoretical work [51, 52]. Its amplitude is normalized, with effective strength absorbed into the coupling coefficients.

The homeostatic variable *H* grows as a saturating function of MA firing rate during wake and decays exponentially during sleep, with state-dependent switching enforced by the Heaviside function *H* (· ) and the threshold potential *V*_*th*_. The coupling *ν*_*vh*_ *>* 0 quantifies the excitatory effect of homeostatic pressure on VLPO recruitment [6, 53], and *τ*_*hm*_ and *τ*_*hv*_ govern, respectively, the buildup and dissipation of H [54]. The constant inputs *A*_*m*_, *A*_*v*_, and *A*_*x*_ aggregate time-averaged drives from sources not modeled explicitly here, e.g., cholinergic input to MA, residual excitatory inputs to Orx, and constant offsets of the circadian drive. Nominal parameter values follow the PR model [6], with additional couplings introduced for the Orx population; all definitions and values are summarized in Table 1.

### 4.3 Analytical derivation of sleep-wake transition thresholds

Here we derive sleep-wake transition thresholds directly from the steady states of the wake-active and sleep-active neural populations in the fast subsystem of the model. Exploiting the strong timescale separation 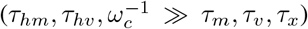, the wake- and sleep-promoting populations can be treated as fast variables responding quasi-statically to the slowly varying net drives (*D*_*v*_, *D*_*x*_). This yields the reduced fast subsystem:

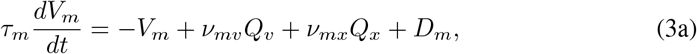

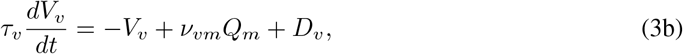

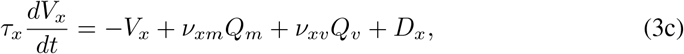

where *D*_*m*_ = *A*_*m*_, *D*_*v*_ = *ν*_*vc*_*C* + *ν*_*vh*_*H* + *A*_*v*_, and *D*_*x*_ = *ν*_*xc*_*C* + *A*_*x*_. Now, sleep-wake transition thresholds can be derived directly from the steady states of the fast neural populations.

At equilibrium, the mean membrane potentials of the wake-active MA population, the sleep-active VLPO population, and the x population satisfy the self-consistency relations,

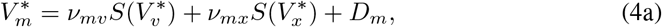

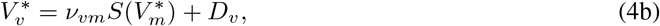

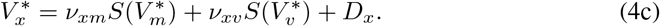

The Jacobian matrix **J** at the equilibrium point 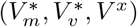 characterizes the local flow, is obtained by taking the partial derivatives of the dynamical equations with respect to the state variables:

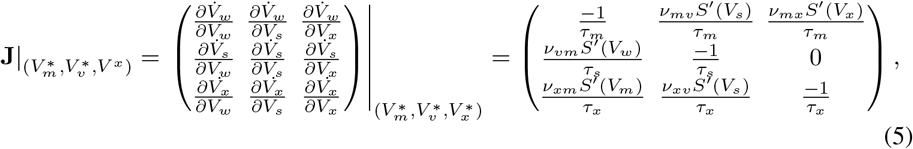

With

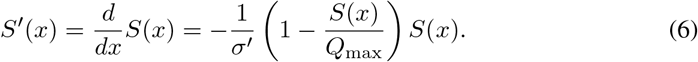

Finally, we arrive at the characteristic equation:

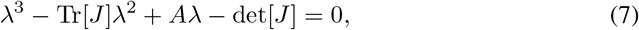

where Tr[*J*] is the trace of the Jacobian, *A* = *JJ*_11_ + *JJ*_22_ + *JJ*_33_, *JJ*_*ii*_ is the algebraic co-factor of the diagonal element *J*_*ii*_, and det[*J*] is the determinant of the Jacobian matrix. The eigenvalues can be given as

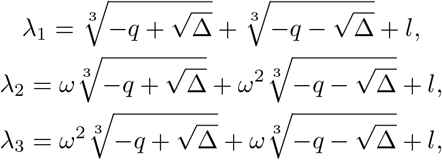

where

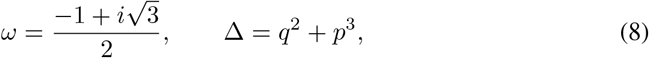

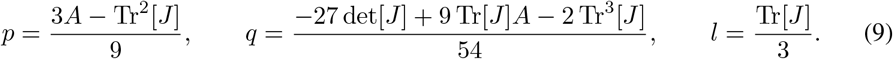

The local stability of the equilibrium state is determined by the real parts of these eigenvalues. In particular, the equilibrium is locally stable if all eigenvalues have negative real parts, whereas a loss of stability occurs when at least one eigenvalue crosses the imaginary axis.

This procedure yields two analytical threshold curves that relate the effective sleep drive *D*_*s*_ to the wake drive *D*_*x*_,

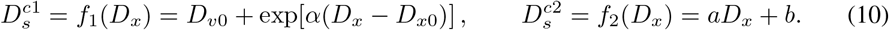

where *α, β, a, b, D*_*v*0_, *D*_*x*0_ are fitting coefficients. *f*_1_ and *f*_2_ are shown in Fig. 1b. The upper boundary, corresponding to sleep onset, is well fitted by an exponential function, whereas the lower boundary, corresponding to awakening, is approximately linear.

Substituting the circadian and homeostatic components into *D*_*v*_ and *D*_*x*_ gives explicit time-dependent thresholds for sleep onset and awakening:

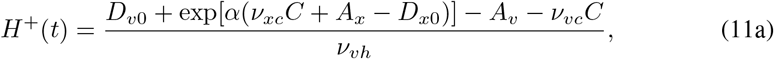

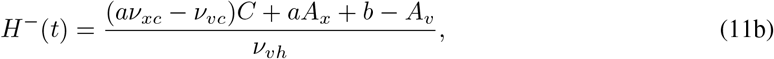

representing the amount of homeostatic sleep pressure required to trigger sleep onset (*H*^+^) and awakening (*H*^−^) at given circadian phase. In the model, sleep occurs when *H*(*t*) exceeds *H*^+^(*t*), and wake occurs when *H*(*t*) falls below *H*^−^(*t*).

## Acknowledgments

This work was supported partially by the Humanities and Social Science Fund of Ministry of Education of China under Grant Nos. 25YJAZH212, the Natural Science Foundation of Zhejiang Province under Grant Nos. LY24A050003, and Jiaxing Public Welfare Project under No. 2025CGZ037.

## Competing interests

All authors declare they have no competing interests.

## Notes

### Competing Interest Statement

The authors have declared no competing interest.

## References

[1] Borbeély, A. A two process model of sleep regulation. Human neurobiology 1, 195—204 (1982).

[2] Daan, S., Beersma, D. G. & Borbely, A. A. Timing of human sleep: recovery process gated by a circadian pacemaker. Am J Physiol 246, R161–83 (1984).

[3] Borbely, A. A. & Achermann, P. Sleep homeostasis and models of sleep regulation. J Biol Rhythms 14, 557–568 (1999).

[4] Borbeély, A. A., Daan, S., Wirz-Justice, A. & Deboer, T. The two-process model of sleep regulation: a reappraisal. J Sleep Res 25, 131–43 (2016).

[5] Phillips, A. & Robinson, P. A quantitative model of sleep-wake dynamics based on the physiology of the brainstem ascending arousal system. Journal of Biological Rhythms 22, 167–179 (2007).

[6] Phillips, A. J. K. & Robinson, P. A. Sleep deprivation in a quantitative physiologically based model of the ascending arousal system. J Theor Biol 255, 413–423 (2008).

[7] Skeldon, A. C.Dijk, D.-J. & Derks, G. Mathematical models for sleep-wake dynamics: Comparison of the two-process model and a mutual inhibition neuronal model. PLOS ONE 9, 1–16 (2014).

[8] Yang, D.-P., McKenzie-Sell, L., Karanjai, A. & Robinson, P. A. Wake-sleep transition as a noisy bifurcation. Phys. Rev. E 94, 022412 (2016).

[9] Sakurai, T. The neural circuit of orexin (hypocretin): maintaining sleep and wakefulness. Nature Reviews Neuroscience 8, 171–181 (2007).

[10] Saper, C. B., Fuller, P. M., Pedersen, N. P., Lu, J. & Scammell, T. E. Sleep state switching. Neuron 68, 1023–1042 (2010).

[11] Chemelli, R. M., Willie, J. T., Sinton, C. M. et al. Narcolepsy in orexin knockout mice: molecular genetics of sleep regulation. Cell 98, 437–451 (1999).

[12] Peyron, C., Faraco, J., Rogers, W. et al. A mutation in a case of early onset narcolepsy and a generalized absence of hypocretin peptides in human narcoleptic brains. Nature Medicine 6, 991–997 (2000).

[13] Thannickal, T. C., Moore, R. Y., Nienhuis, R. et al. Reduced number of hypocretin neurons in human narcolepsy. Neuron 27, 469–474 (2000).

[14] Fulcher, B. D., Phillips, A. J. K., Postnova, S. & Robinson, P. A. A physiologically based model of orexinergic stabilization of sleep and wake. PLOS ONE 9, 1–14 (2014).

[15] Li, S.-B. et al. Hyperexcitable arousal circuits drive sleep instability during aging. Science 375, eabh3021 (2022).

[16] Mander, B. A., Winer, J. R. & Walker, M. P. Sleep and human aging. Neuron 94, 19–36 (2017).

[17] Van Cauter, E., Leproult, R. & Plat, L. Age-related changes in slow wave sleep and rem sleep and relationship with growth hormone and cortisol levels in healthy men. JAMA 284, 861–868 (2000).

[18] Robinson, P. A., Phillips, A. J. K., Fulcher, B. D., Puckeridge, M. & Roberts, J. A. Quantitative modelling of sleep dynamics. Philos Trans A Math Phys Eng Sci 369, 3840–54 (2011).

[19] Saper, C. B., Chou, T. C. & Scammell, T. E. The sleep switch: hypothalamic control of sleep and wakefulness. Trends Neurosci 24, 726–731 (2001).

[20] Saper, C. B., Scammell, T. E. & Lu, J. Hypothalamic regulation of sleep and circadian rhythms. Nature 437, 1257 EP – (2005).

[21] Gallopin, T. et al. Identification of sleep-promoting neurons in vitro. Nature 404, 992–5 (2000).

[22] Lu, J., Greco, M. A., Shiromani, P. & Saper, C. B. Effect of lesions of the ventrolateral preoptic nucleus on nrem and rem sleep. J Neurosci 20, 3830–3842 (2000).

[23] Hagan, J. J. et al. Orexin A activates locus coeruleus cell firing and increases arousal in the rat. Proceedings of the National Academy of Sciences USA 96, 10911–10916 (1999).

[24] Yamanaka, A. et al. Orexins activate histaminergic neurons via the orexin 2 receptor. Biochemical and Biophysical Research Communications 290, 1237–1245 (2002).

[25] Liu, R.-J., van den Pol, A. N. & Aghajanian, G. K. Hypocretins (orexins) regulate serotonin neurons in the dorsal raphe nucleus by excitatory direct and inhibitory indirect actions. The Journal of Neuroscience 22, 9453–9464 (2002).

[26] Yamanaka, A., Muraki, Y., Tsujino, N., Goto, K. & Sakurai, T. Regulation of orexin neurons by the monoaminergic and cholinergic systems. Biochemical and Biophysical Research Communications 303, 120–129 (2003).

[27] Muraki, Y. et al. Serotonergic regulation of the orexin/hypocretin neurons through the 5-HT1A receptor. The Journal of Neuroscience 24, 7159–7166 (2004).

[28] Li, Y. & van den Pol, A. N. Direct and indirect inhibition by catecholamines of hypocretin/orexin neurons. The Journal of Neuroscience 25, 173–183 (2005).

[29] Xie, X. et al. GABAB receptor-mediated modulation of hypocretin/orexin neurones in mouse hypothalamus. The Journal of Physiology 574, 399–414 (2006).

[30] Matsuki, T. et al. Selective loss of GABA_B_ receptors in orexin-producing neurons results in disrupted sleep/wakefulness architecture. Proceedings of the National Academy of Sciences USA 106, 4459–4464 (2009).

[31] Colwell, C. S. Linking neural activity and molecular oscillations in the scn. Nature Reviews Neuroscience 12, 553–569 (2011).

[32] Hastings, M. H., Maywood, E. S. & Brancaccio, M. Generation of circadian rhythms in the suprachiasmatic nucleus. Nature Reviews Neuroscience 19, 453–469 (2018).

[33] Aston-Jones, G., Chen, S., Zhu, Y. & Oshinsky, M. L. A neural circuit for circadian regulation of arousal. Nature Neuroscience 4, 732–738 (2001).

[34] Porkka-Heiskanen, T. et al. Adenosine: A mediator of the sleep-inducing effects of prolonged wakefulness. Science 276, 1265–1268 (1997).

[35] Morairty, S., Rainnie, D., McCarley, R. & Greene, R. Disinhibition of ventrolateral preoptic area sleep-active neurons by adenosine: a new mechanism for sleep promotion. Neuroscience 123, 451–457 (2004).

[36] Urade, Y. & Hayaishi, O. Prostaglandin d2 and sleep regulation. Biochimica et Biophysica Acta (BBA) - Molecular and Cell Biology of Lipids 1436, 606–615 (1999).

[37] Kumar, S. et al. Adenosine a(2a) receptors regulate the activity of sleep regulatory gabaergic neurons in the preoptic hypothalamus. Am J Physiol Regul Integr Comp Physiol 305, R31–41 (2013).

[38] Yao, C., Jiang, J., Gu, C., Shuai, J. & Yang, D. Circadian-modulated thresholds as a mechanistic basis for sleep-wake transitions, recovery, and sleepiness. bioRxiv 2026.02.04.703688 (2026).

[39] Athanasouli, C., Piltz, S. H., Diniz Behn, C. G. & Booth, V. Bifurcations of sleep patterns due to homeostatic and circadian variation in a sleep-wake flip-flop model. SIAM Journal on Applied Dynamical Systems 21, 1893–1929 (2022).

[40] Pizza, F. et al. Clinical and polysomnographic course of childhood narcolepsy with cataplexy. Brain 136, 3787–3795 (2013).

[41] Hong, J. et al. Personalized sleep-wake patterns aligned with circadian rhythm relieve daytime sleepiness. iScience 24 (2021).

[42] Song, Y. M. et al. A real-time, personalized sleep intervention using mathematical modeling and wearable devices. Sleep 46, zsad179 (2023).

[43] Skeldon, A. C. et al. Method to determine whether sleep phenotypes are driven by endogenous circadian rhythms or environmental light by combining longitudinal data and personalised mathematical models. PLOS Computational Biology 19, 1–33 (2023).

[44] Shochat, T., Santhi, N., Herer, P.Dijk, D.-J. & Skeldon, A. C. Sleepiness is a signal to go to bed: data and model simulations. Sleep 44, zsab123 (2021).

[45] Diniz Behn, C. G. & Booth, V. Simulating microinjection experiments in a novel model of the rat sleep–wake regulatory network. Journal of Neurophysiology 103, 1937–1953 (2010).

[46] Ji, Y., Xu, F., Shuai, J., Yang, D. & Yao, C. Dynamical mechanism for the interplay of circadian, homeostatic, and ultradian rhythm in normal human sleep. Phys. Rev. E 111, 044215 (2025).

[47] Herring, W. J. et al. Orexin receptor antagonism for treatment of insomnia: a randomized clinical trial of suvorexant. Neurology 79, 2265–2274 (2012).

[48] Forger, D. B., Jewett, M. E. & Kronauer, R. E. A simpler model of the human circadian pacemaker. Journal of Biological Rhythms 14, 532–537 (1999).

[49] Gooley, J. J., Schomer, A. & Saper, C. B. The dorsomedial hypothalamic nucleus is critical for the expression of food-entrainable circadian rhythms. Nat Neurosci 9, 398–407 (2006).

[50] Chou, T. Regulation of Sleep-wake Timing: Circadian Rhythms and Bistability of Sleep-wake States (Harvard University, 2003).

[51] Aschoff, J. Circadian rhythms in man. Science 148, 1427–1432 (1965).

[52] Gu, C., Xu, J., Liu, Z. & Rohling, J. H. T. Entrainment range of nonidentical circadian oscillators by a light-dark cycle. Phys. Rev. E 88, 022702 (2013).

[53] Phillips, A. J. K., Fulcher, B. D., Robinson, P. A. & Klerman, E. B. Mammalian rest/activity patterns explained by physiologically based modeling. PLoS Comput Biol 9, e1003213 (2013).

[54] Rusterholz, T., Duürr, R. & Achermann, P. Inter-individual differences in the dynamics of sleep homeostasis. Sleep 33, 491–498 (2010).

